# High throughput functional screening for next generation cancer immunotherapy using droplet-based microfluidics

**DOI:** 10.1101/2020.11.25.399188

**Authors:** Yuan Wang, Ruina Jin, Bingqing Shen, Na Li, He Zhou, Wei Wang, Yingjie Zhao, Mengshi Huang, Pan Fang, Shanshan Wang, Pascaline Mary, Ruikun Wang, Peixiang Ma, Ruonan Li, Youjia Cao, Fubin Li, Liang Schweizer, Hongkai Zhang

## Abstract

Currently high throughput approaches are lagged for isolation of antibodies whose function goes beyond simple binding, which have prevented the next generation cancer immunotherapeutics, such as bispecific T cell engager antibodies or agonist antibody of costimulatory receptor, from reaching their full potential. Here we developed a highly efficient droplet-based microfluidics platform combining with lentivirus transduction system that enables functional screening of millions of antibodies. To showcase the capacity of the system, functional antibodies for CD40 agonism with low frequency (<0.02%) were identified with 2 rounds of screening. To demonstrate its versatility, an anti-Her2/anti-CD3 bispecific antibody library was established using bispecific T cell Engager (BiTE) platform and functional screening enabled efficient identification of potent anti-Her2/anti-CD3 BiTE antibodies. The platform could revolutionize the next generation cancer immunotherapy drug development and research world.

## INTRODUCTION

Cancer immunotherapies harness the power of the immune system to treat tumor and has brought huge success and driven a major paradigm shift in cancer treatment. The first generations of cancer immunotherapy agents consist primarily of antagonist antibodies that block negative immune checkpoints, such as programmed cell death protein 1 (PD-1) *(1–3)*. Recently, the next generation cancer immunotherapies, including bi/multi-specific antibodies such as bispecific T-cell engager (BiTE) antibodies and agonist antibodies of costimulatory receptors are getting into the spotlight and showed promises in anticancer immunity *(4–7)*.

The in vitro display technology, such as phage display, allows to select antibody binders from large combinatorial library, with the capacity of 10^11^ diversities *(8–13)* and has been one of the major players in conventional antibody drug development by the use of high or low throughput simple binding assays.

Next generation cancer immunotherapies including agonist or bispecific/multi-specific antibodies have yet to fulfill their expectations. This may be, at least in part, due to the low throughput (maximal of a few thousand antibodies can be tested) of functional screens, a bottle neck that requires prescreens to narrow down potential candidates, and may cause the loss of certain target profiles depending on criteria set for the pre-screens. In addition, bi/multi-specific antibody development may also suffer from lack of diversity driven by limited candidate generations using validated monospecific antibodies with known bispecific scaffolds, and largely neglects the compatibilities of different arms of the bi/multi-specific antibodies.

The droplet microfluidics technology allows screening of antibody secreting cells at single cell level that could not be obtained using the bulk population-based assays with an unprecedented throughput. The microfluidics droplet system can encapsulate single cells in the water-in-oil droplets at the rate of thousands of droplets per second *(14)*. Antibodies generated by the cells are contained in the droplet, enabling the maintenance of phenotype and genotype linkage in the droplet *(15–17)*. Finally, the droplets containing desirable cells are sorted by fluorescence activated droplet sorting (FADS). Bachir et al described application of microfluidics droplet system to screen hundred-thousands of hybridoma cells for antibodies that inhibit enzyme ACE-1 or bind to target cells *(18, 19)*. With the sophistication of the microfluidics droplet system, simultaneous analyses of millions of individual antibody secreting plasma cells were achieved for their antibody secretion rate and affinity *(20)*. And millions of plasma cells were screened for antibodies bound to vaccine or cancer target *(21)*. However, improvement is in urgent need to screen for functional antibodies.

Here an efficient technology platform was developed to simultaneously screen the binding and the agonistic activity of antibodies or the functions of bispecific antibodies by combining the strength of an autocrine based lentivirus transduction system developed by Hongkai et al. *(22, 23)* with microfluidics droplet system. The technical capabilities of the platform were demonstrated by the successful identification of rare potent costimulatory receptor CD40 agonist antibodies (0.02%) and anti-Her2/anti-CD3 bispecific antibodies from combinatorial antibody library. The streamlined technology enables the efficient and unbiased discovery of active antibodies for the next generation of cancer immunotherapy with dramatically increased throughput and accuracy.

## RESULTS

### Outline of function-based antibody selection using droplet-based microfluidics

We designed the functional antibody selection platform as depicted in Fig. 1. Cells were infected by the lentiviral antibody expressing library at multiplicity of infection (MOI) equal to 1 so that each cell express and secrete only one type of antibody. For microfluidics system based screening, individual antibody secreting cell was coencapsulated with a reporter cell into a droplet by microfluidic dropmaker. The resulting emulsion was incubated off-chip overnight and injected into the sorting chip. The droplets containing activated reporter cells were sorted by fluorescence activated droplet sorting (FADS). Subsequently, the cells were recovered from the droplets and the functional antibodies were identified by sequencing the antibody genes in the sorted cells.

**Fig. 1.**
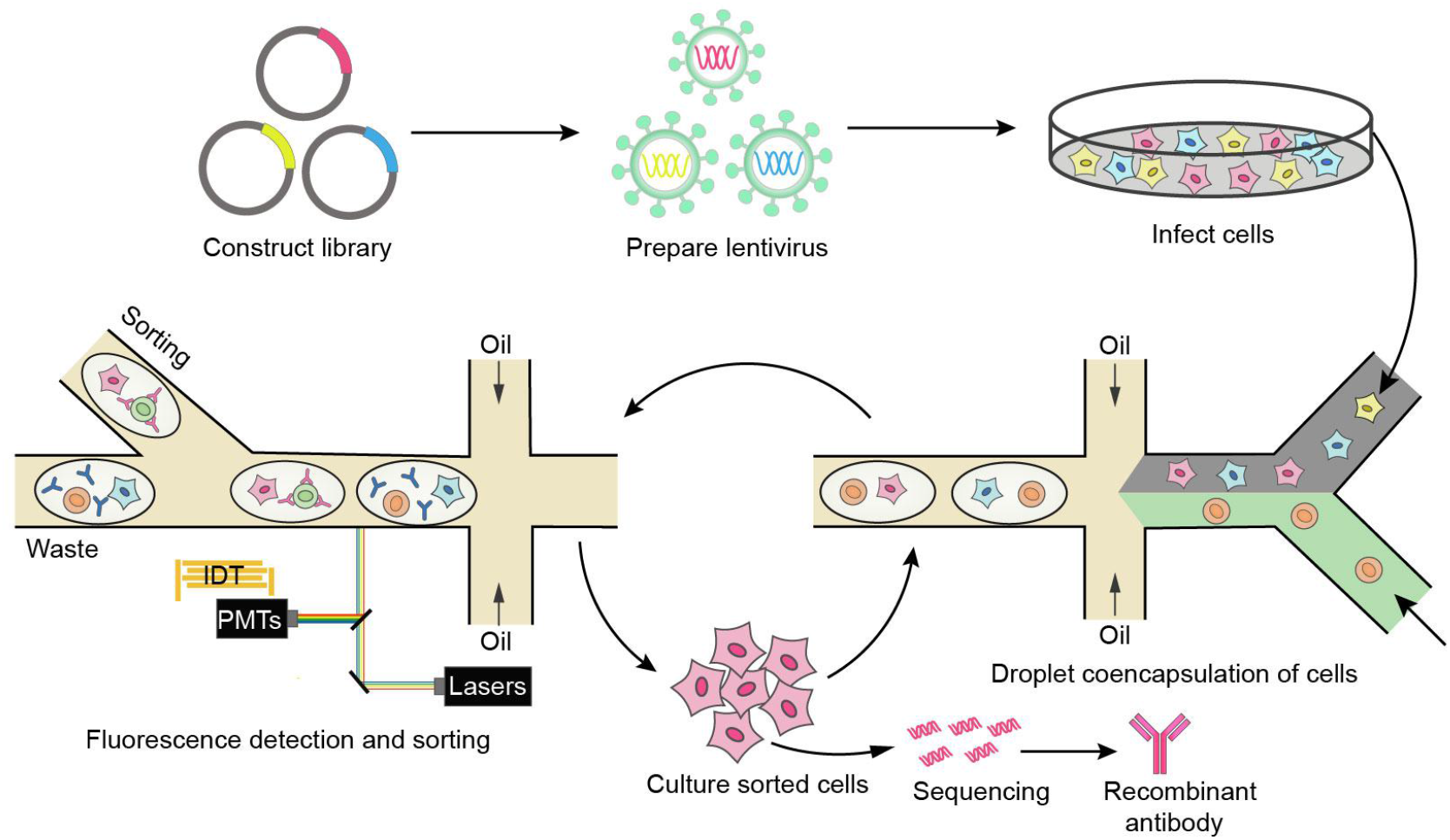
Schematic overview of functional screening antibody by droplet-based microfluidics. The antibody genes were cloned into lentiviral vectors. Eukaryotic cells were infected by the lentiviral antibody library and individual transduced cells were coencapsulated with the reporter cell into droplets using microfluidics system. The resulting emulsion was incubated off-chip overnight and injected into the sorting chip. Droplets containing antibody secreting cells and activated reporter cell were sorted. The sorted cells were cultured for the next round of selection. After multiple rounds of iteration, antibody genes were amplified from the sorted cells and analyzed by Sanger Sequencing or Next Generation Sequencing. The enriched antibodies were synthesized and recombinant antibodies were expressed and tested for function.

### Development and optimization of droplet based assay

Two microfluidics devices were used (i) to compartmentalize the lentivirus infected cells with the reporter cells and detection reagents (Figure 2A,2B) (ii) to sort droplets based on reporter cell activation and receptor binding signals using surface acoustic wave based sorter (Fig. 2C) *(24)*.

**Fig. 2.**
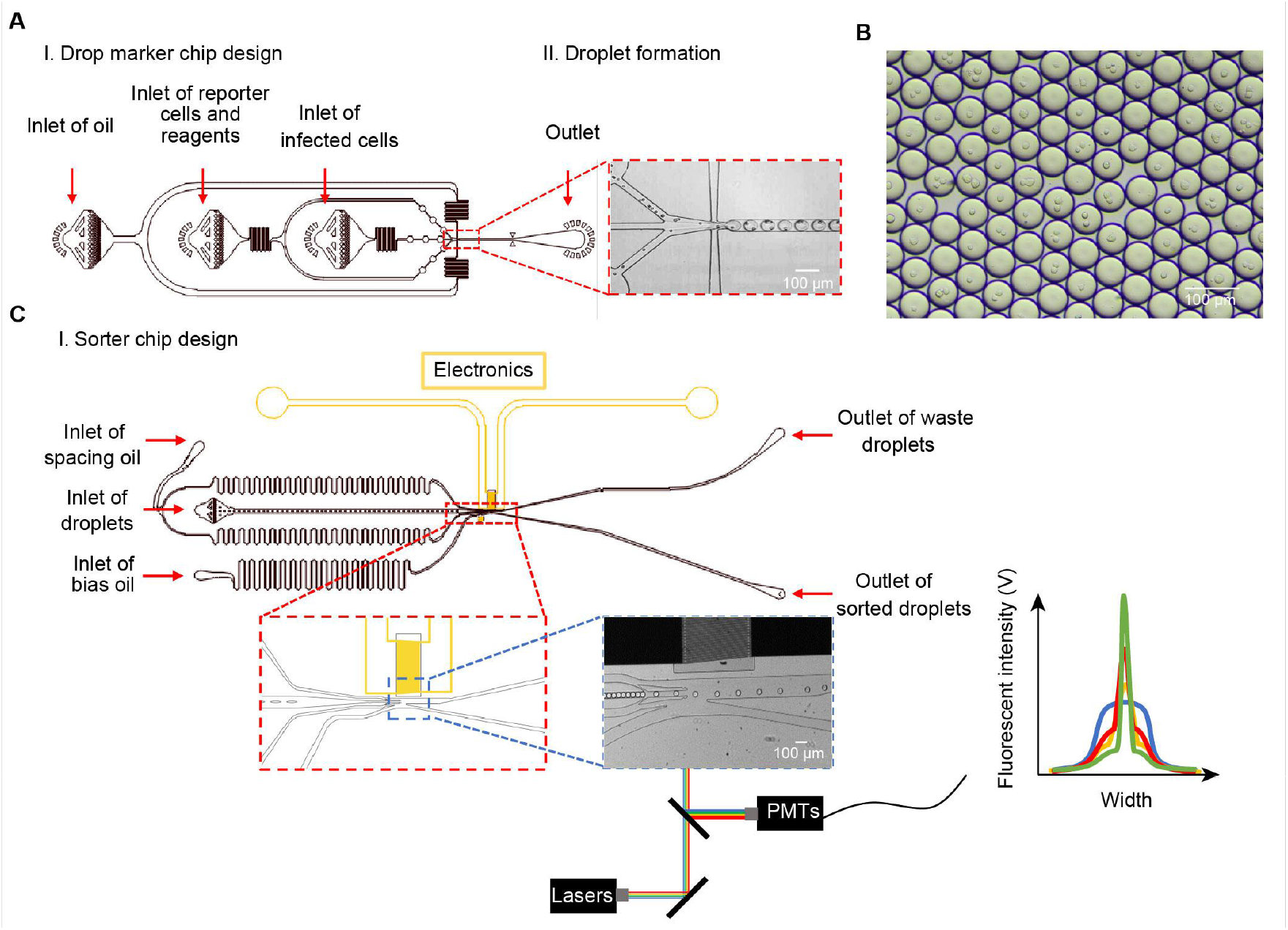
Microfluidics chips. (**A**) The droplet maker chip is used to generate droplets to coencapsulate antibody secreting cells with reporter cells. (**B**) Image of generated droplets. (**C**) The sorting chip is used to collect droplets based on the intensity of fluorescence. The functions of various inlets is indicated and pictures of outlets for droplet generation and sorting were showed.

The droplets generated by microfluidics chip were incubated for 24 h at 37°C. No merging of droplets was observed, indicating that the droplets are stable (data not shown). Next, we used anti-Her2/anti-CD3 BiTE antibody as an example for the development and optimization of the droplet-based assay. Anti-Her2/anti-CD3 positive control was engineered to recognize Her2 on tumor cells and CD3 on T cells using the variable domain sequences of Trastuzumab and blinatumomab, respectively *(25)*. Her2 overexpressing K562-Her2 cells were infected with the anti-Her2/anti-CD3 positive control lentivirus at a low MOI and less than 10% of cells were infected. The K562-Her2 played a dual role to express antibody and provide Her2 mediated crosslinking of the secreted antibodies. Each infected K562-Her2 cells were encapsulated with a Jurkat/pIL2-eGFP reporter cell (fig. S1) with on average 0.5 antibody secreting cell per droplet on average. The number of cells per droplet was analyzed based on the image and the results are in good agreement with double Poisson distribution (fig. S2). After 16h incubation, the droplets were re-injected to the sort chip to analyze the activation of reporter cell in the droplet. About 9.5% reporter cells were activated. In contrast, when the anti-Her2/anti-CD3 lentivirus infected K562-Her2 cells were cocultured with the reporter cells in the plate well, 72.7% of the reporter cells were activated after 16h incubation (Fig. 3A). The different percentage of reporter cell activation demonstrated that antibody secreted by cells in the droplet can’t diffuse between different droplets thus the system can be used for screening of library. In addition, NucGreen Dead 488 were used to track the cell viability in droplets. Both K562 and Jurkat cells in droplets have high viability around 90% after 16h incubation (fig. S3).

**Fig. 3.**
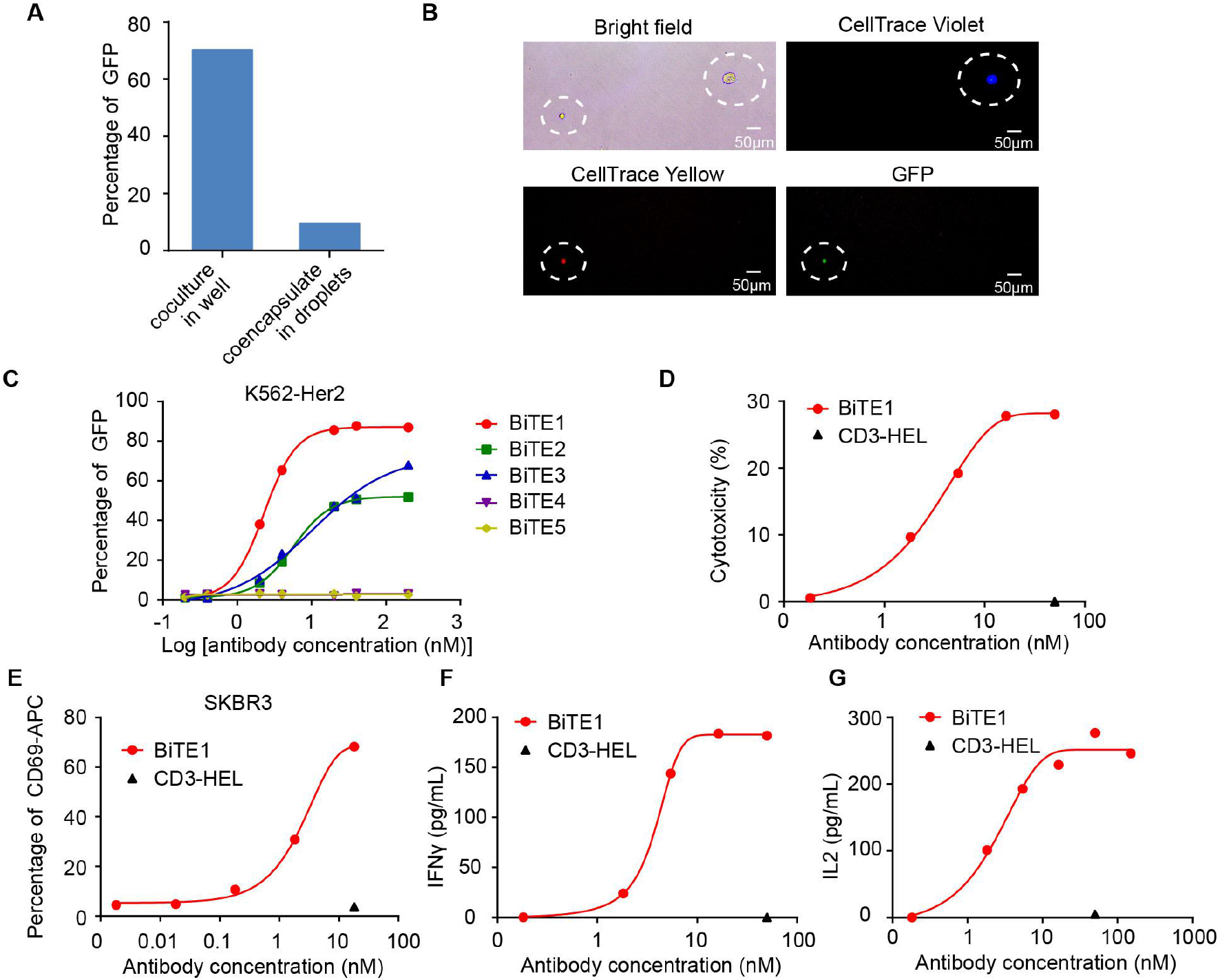
Screening anti-Her2/anti-CD3 BiTE antibody from a bispecific antibody library. (**A**) Activation of reporter cell in the plate-well based or droplet based coculture. The K562-Her2 cells expressing anti-Her2/anti-CD3 BiTE antibody were cocultured with the Jurkat/pIL2-eGFP reporter cell in plate-well or individually coencapsulated with the reporter cell. The activation of reporter cells in both conditions were compared. (**B**) Representative image of a sorted droplet. The antibody secreting K562-Her2 cells and the reporter cells were stained with CellTrace Yellow and CellTrace Violet, respectively. K562-Her2 cells were individually coencapsulated with the reporter cell and the droplets containing activated reporter cell were sorted. (**C**) Activation of reporter cell by the identified antibodies. The K562-Her2 cells were cocultured with the reporter cells in presence of the identified antibodies. Expression of GFP by the reporter cell was analyzed. (**D-G**) PBMC and tumor cell coculture assay. PBMCs and SK-BR-3 cells were cocultured in presence of BiTE1 or control antibody. Tumor cell lysis was determined by measuring the release of Lactic Acid Dehydrogenase (LDH) from tumor cells. (**D**) The activatoin marker CD69 expression on T cells was investigated (**E**) and the levels of cell secreted IFN-γ (**F**) and IL-2 (**G**) in the culture supernatant were measured by ELISA.

### Screening anti-Her2/anti-CD3 bispecific antibody using the function-based screening method

Because the phage display technology can be used to screen antibody library with much larger diversity than is possible in eukaryotic systems, we first selected Her2 binding antibodies from an antibody phage display library. We panned a naïve human single chain Fv (scFv) library with size of 1010 members against Her2 protein. One rounds of panning were performed and the scFv genes in phagemid were subcloned into a lentiviral vector which contained a fixed blinatumomab derived anti-CD3 scFv gene to express anti-Her2×antib-CD3 BiTE antibody. The size of the combinatorial bispecific antibody library is about 10^5^ members.

K562-Her2 cells were infected with the BiTE antibody lentivirus library at low MOI to ensure most cells were infected by only one virus and produced a single type of bispecific antibody. Prior cell encapsulation, K562-Her2 cells and Jurkat/pIL2-eGFP cells were stained with CellTrace Violet and CellTrace Yellow dye,respectively. The infected K562-Her2 cells were individually coencapsulated with a Jurkat/pIL2-eGFP reporter cell into the same droplet. After 16h incubation, the droplets were sorted (fig. S4). We first selected droplets for the presence of K562-Her2 and Jurkat/pIL2-eGFP cells based on the cell staining fluorescence signals, then we selected droplets if the reporter cell inside was activated based on GFP signal and finally we selected the droplets if the signal of GFP was colocalized with the reporter Jurkat cell staining signal. 0.26% droplets were sorted and the image of a representative sorted droplet was shown (Fig. 3B). The anti-Her2 scFv genes were amplified from the sorted cells and cloned into mammalian expression plasmid. Twenty clones were picked for Sanger sequencing and five bispecific antibodies appearing more than once. The purified bispecific antibodies were added into the Jurkat/pIL2-eGFP reporter cell in coculture with K562-Her2. Three out of five antibodies (BiTE-1, BiTE-2 and BiTE-3) can activate the reporter cell in presence of K562-Her2 (Fig 3C). To assess the anti-tumor activity of BiTE1, we performed in vitro cytotoxicity assays by coculturing PBMCs and HER2-expressing SKBR3 cells. The results showed that lysis of SKBR3 cells only occurred when it was incubated with BiTE1, not with control antibodies CD3-HEL (Fig. 3D). Flow cytometry analysis of T cells in the coculture system showed that BiTE1 stimulated the expression of activation marker CD69 on T cells (Fig. 3E). Moreover, BiTE1 induced dose-dependent increases of IFN-γ and IL-2 in the supernatants (Fig. 3F and 3G).

On the whole, the results demonstrated that this technology allows combinatorial screening and profiling of large numbers of bispecific antibodies.

### Proof of concept screening of CD40 agonist from a spike-in library

To further validate the utility of the function-based screening method, we sought to identify CD40 agonistic antibodies. CD40 is a promising drug target for cancer immune therapy *(5)*. Activation of CD40 on antigen presenting cells (APCs) results in improved antigen processing and presentation, and cytokine release, which enhances T cell response. Human Jurkat T cells were engineered to express human CD40 and express GFP controlled by NF-κB response elements. The reporter cell line express GFP when the CD40 is activated (fig. S5). HEK293FT cells coexpressing red fluorescence protein (RFP) and hexameric form of CD40L was used as positive control (CD40L cell). HEK293FT cells coexpressing blue fluorescence protein (BFP) and hen egg lysozyme (HEL) antibody was used as negative control (HEL cell). The CD40L or HEL cells were coencapsulated with the Jurkat/NF-κB-GFP-hCD40 reporter cells into droplets. After 16h incubation, 24% of the droplets containing the CD40L cell and reporter cell exhibited GFP fluorescence signal while the HEL cell and reporter cell coencapsulating droplets showed clean background of activation of the reporter cell (0.5%) (fig. S6).

CD40L cells were spiked into a 10-fold excess of HEL cells. The mixture of CD40L cells and HEL cells were coencapsulated with the Jurkat/NF-κB-GFP-hCD40 reporter cells with 0.5 protein secreting cell per droplet. The droplets which contained activated reporter cell were sorted based on green fluorescence. Before sorting only 1.24% of the mixed cell population were RFP positive, 16.79% of cell were BFP positive and the rest droplets were empty or contained only reporter cell, whereas after the sort the percentage of droplets containing RFP positive CD40L cell increased to 51.94% where 49.24% were RFP positive and 2.70% were RFP and BFP double positive (Fig. 4A). We also observed that most droplets containing both RFP positive CD40L cells and activated GFP reporter cell after sorting (Fig. 4B).

**Fig. 4.**
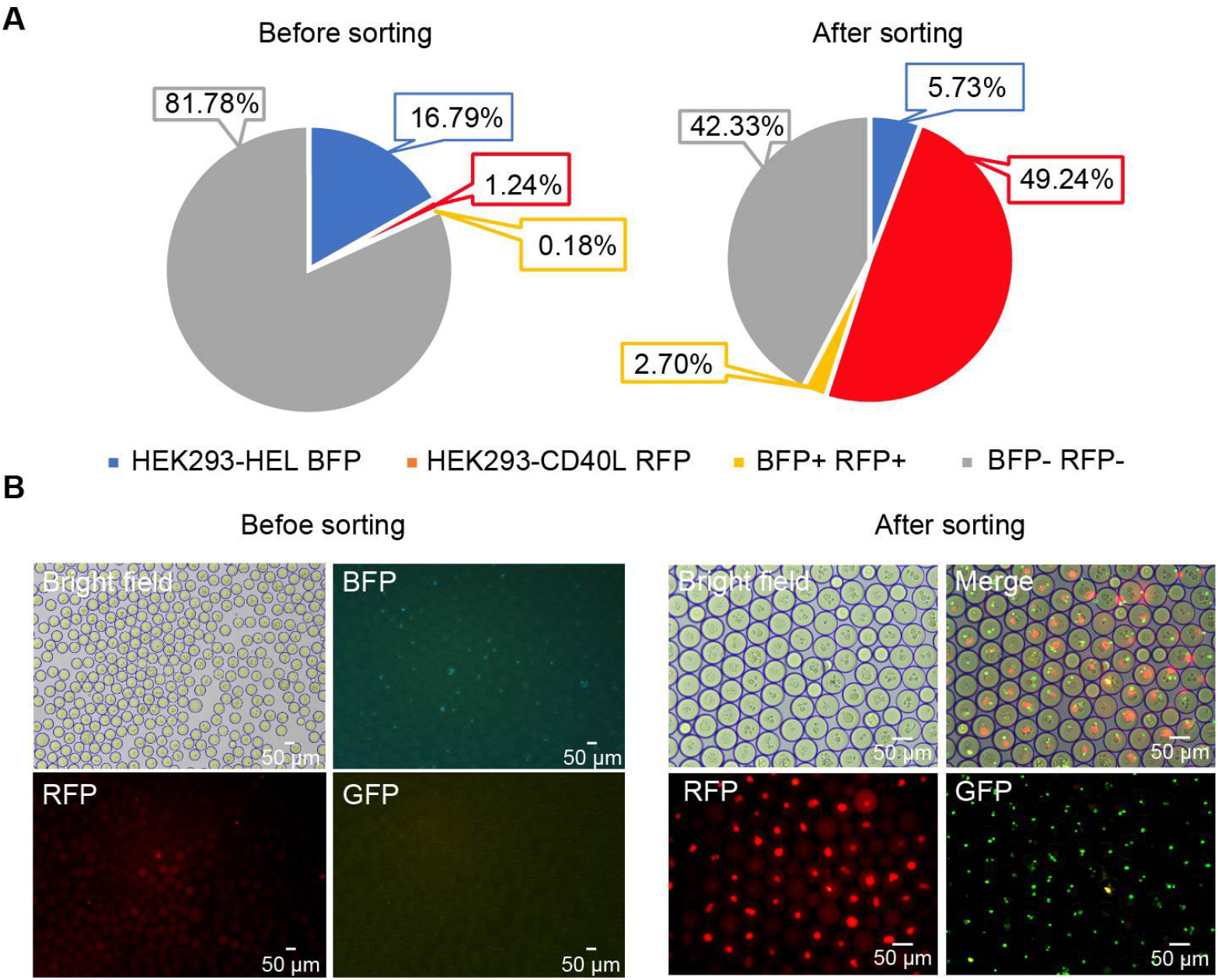
Screening CD40 agonist from a spike-in library with microfluidics system. (**A**) RFP-positive hexameric CD40L protein secreting cells were spiked into a 10-fold excess of BFP-positive nonrelated anti-HEL antibody secreting cells and the mixture of cells were coencapsulated with the reporter cells. After incubation, the droplets containing activated reporter cells were sorted. The proportion of droplets containing RFP-positive or BFP-positive cells before and after sorting was analyzed. (**B**) Bright field and fluorescence images of droplets before and after sorting.

### Screening CD40 agonist antibody using the function-based screening method

We first selected CD40 binding antibodies from antibody library using phage display technology. The scFv genes in phagemid were subcloned into lentiviral vector that contained IgG1 Fc gene to express scFv Fc fusion protein. The library size was about 10^5^. HEK293FT cells were infected with lentivirus library at low MOI to ensure most cells were infected by only one virus and produced one type of monoclonal antibody.

To screen CD40 agonist antibodies, the antibody producing cells were coencapsulated with CellTrace Yellow prestained Jurkat/NF-κB-GFP-hCD40 reporter cell and Dylight647 conjugated secondary antibody in droplets. The reporter cells were also coencapsulated with soluble hexameric CD40L protein and anti-HEL antibody as positive and negative controls. Droplets of different populations (positive control, negative control and the screening population) were coded by adding different concentrations of DY405. After 16h incubation, the droplets were sorted based on the following criteria (fig. S7). The droplets of the screening population were first gated based on the intensity of DY405. CellTrace Yellow fluorescence signal showed the presence of reporter cell in the droplet. The Dylight647 fluorescence signal indicated binding of the secreting antibodies to CD40 on the surface of the reporter cell and GFP fluorescence signal peak indicated activation of the reporter cell (fig. S7). The droplets displayed different patterns of fluorescence signals (Fig. 5A, fig. S8). Whereas only 0.29% droplets were Dylight647 and green fluorescence double positive for the 1^st^ round of screening (out of the droplet containing Jurkat cell signal), this value increased to 12.5% for the 2^nd^ round of screening (Fig. 5B). The droplets after sorting were imaged and the number of Dylight647 and green fluorescence double positive droplets dramatically increased after the sorting (Fig. 5C).

**Fig. 5.**
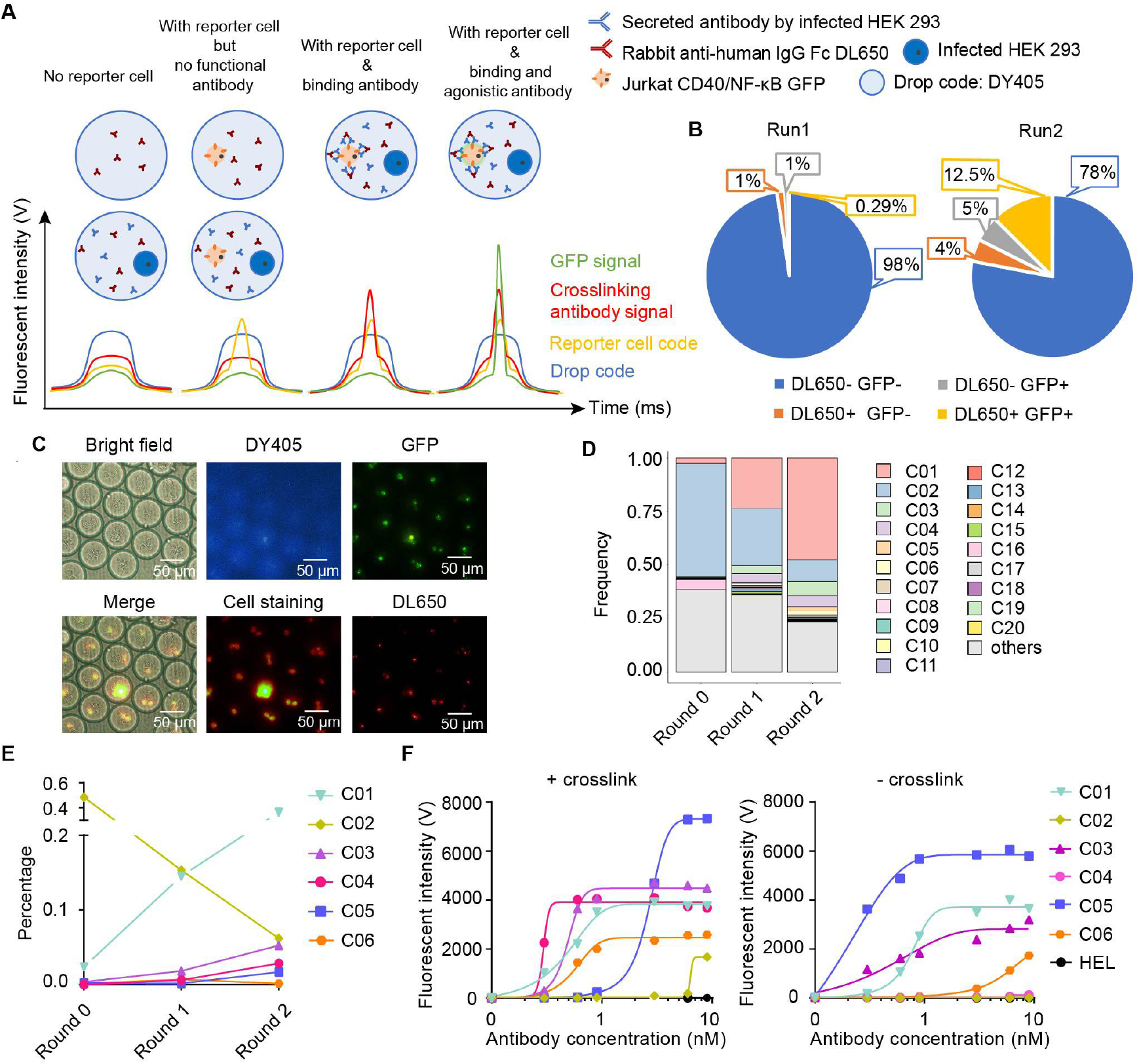
Screening CD40 agonist antibody from monoclonal antibody library. HEK293T cels were infected with a lentivirus antibody library and individually coencapsulated with Jurkat reporter cell and fluorescence labeled secondary antibody in droplets. The droplets containing reporter cell activated by antibodies secreted by the coencapsulated antibody expressing cell were sorted. The sorted cells were expanded for the 2^nd^ round of selection. The enriched antibodies were identified by NGS. (**A**) Schematic of possible time traces. (**B**) The proportions of different types of droplets for each round of selection was analyzed. (**C**) Bright field and fluorescence images of the sorted droplets after the second round of selection. Bar plot for the top 20 scFv clusters and their frequencies during the selection process. The change of frequencies of the selected antibodies during the selection process. (**F**) Agonist activity of the selected antibodies was determined using the CD40 reporter cell line in presence or absence of crosslinking secondary antibody.

The scFv genes were amplified from the cells after each round of sorting and were subject to the third-generation sequencing. Circular consensus sequences from these reads were generated, filtered by quality and full-length scFvs were identified in 44084 reads. Considering the errors introduced by PCR or sequencing, the similar (at 95% similarity) scFv sequences were grouped into 561 scFv clusters and consensus scFv sequences of each cluster were created. Comparison the sequence frequency in each round revealed some scFvs were enriched while some scFvs were eliminated during the selection (Fig. 5D). The frequency of scFv clusters C01, C03, C04, C05 and C06 were low before sorting, and showed roundwise increase during the selection process. In contrast, scFv cluster C02 were highly abundant before sorting and its frequency was significantly reduced (Fig. 5E).

The genes of full length IgG C01, C02, C03, C04, C05 and C06 were synthesized and recombinant antibodies were expressed and purified. The activity of these antibodies was tested using the Jurkat/NF-κB-GFP-hCD40 reporter cell line. The reporter cells were stimulated with different concentrations of antibody in the presence or absence of the crosslinking secondary antibody. Antibodies C01, C03, C04, C05 and C06 can activate the reporter cell line. The activity of C01, C03 and C05 was independent of crosslinking while C04 and C06 activated the reporter cell in a crosslinking dependent manner (Fig. 5F). The Fc receptor mediated crosslinking independent activity of CD40 agonist antibody is concerned because this feature underlied the systemic adverse events. Therefore, the crosslinking dependent antibody C04 was chosen for further characterization.

Flow cytometry and SPR results showed that antibody C04 bound to human, rhesus macaque and cynomolgus monkey CD40 with similar affinity (fig. S9A and S9B). In addition, ELISA results confirmed that antibody C04 specifically bound to CD40 rather than other TNF receptor superfamily members such as GITR, OX40 and 4-1BB (fig. S9C). Given that FcγRIIB expressed on tumor infiltrating myeloid cells is required for agonistic activity of the crosslinking dependent CD40 agonist antibodies, the FcγRIIB dependency of antibody C04 was confirmed. The Jurkat/NF-κB-GFP-hCD40 reporter can be stimulated by C04 in the presence of FcγRIIB-overexpressing cells. The results showed that the agonism of C04 was FcγRIIb dependent (Fig. 6A).

**Fig. 6.**
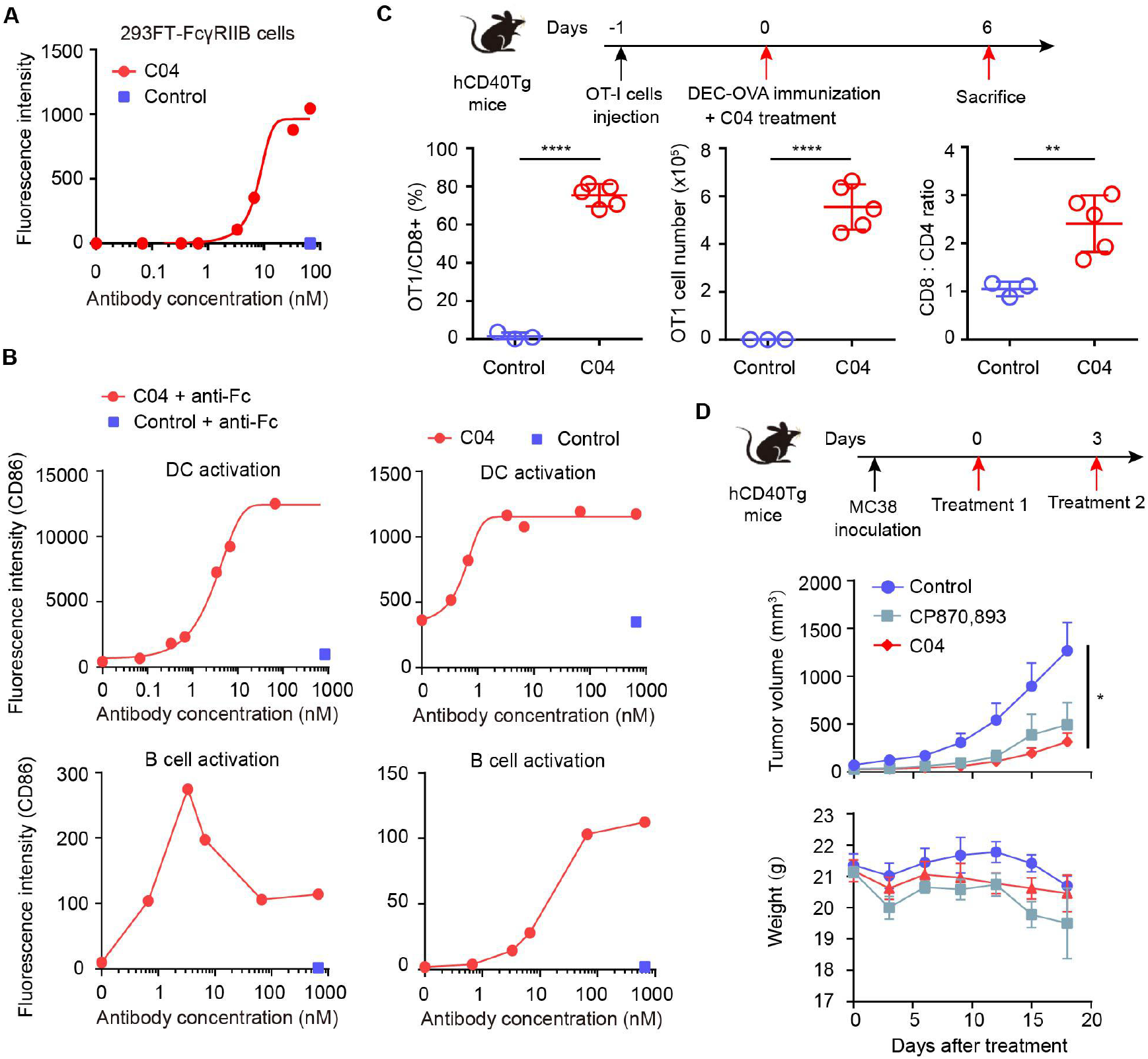
Characterization of antibody in in vitro and in vivo models. (A) The FcγRIIB dependency of antibody C04. Jurkat/NF-κB-GFP-hCD40 reporter cells were stimulated by antibody C04 or anti-HEL control in the presence of FcγRIIB overexpressing HEK293T cells. The activation of reporter cell line were analyzed by flow cytometry. (B) The activation of DCs or B cells by C04. Human DC cells or B cells were stimulated by C04 in the absence or presence of anti-Fc antibody. The expression of CD86 was analyzed by flow cytometry. (C) OVA-specific CD8+ T cell response induced by C04 in the CD40/FcgR humanized mice. The transgenic mice were adoptively transfered with OVA-specific OT-I cells and treated with DEC-OVA together with C04 or isotype control antibody. Mice were euthanized for the analysis of T cells. Each circle represented an individual mouse. (D) Antitumor effect of C04 in syngeneic mouse model. CD40/FcgR humanized mice were s.c. engrafted with MC38 tumor cells. When MC38 tumors were established (∼100 mm^3^), mice were treated with C04, CP-870,893 or isotype control antibody. The tumor volume and body weight were measured every three days until the end of the experiment. Data are represented as mean ± SEM.

To investigate whether C04 can promote the activation of CD40-positive APCs, human dendritic cells (DCs) and B cells isolated from PBMC were stimulated by C04 in the presence or absence of the crosslinking antibody. Flow cytometry analysis showed that C04 upregulated the expression of activation marker CD86 on DCs and B cells and anti-Fc mediated crosslinking could further enhance the agonistic activity (Fig. 6B).

The immunostimulatory activity of antibody C04 was further assessed by OVA-specific CD8+ T cell response model in FcγR/CD40-humanized mice. Mice treated with CD40 agonist antibody showed an increased number and percentage of OT-I cells among CD8+ T cells, as compared to mice treated with the isotype antibody, demonstrating C04 had robust adjuvant effect (Fig. 6C).

In addition, antitumor efficacy of C04 was assessed in syngeneic tumor models. When MC38 tumors were established (∼100 mm^3^), mice were treated with C04, CP-870,893 (Pfizer) or anti-HEL antibodies. C04 displayed comparable anti-tumor activity as CP-870,893, which is the most potent CD40 agonistic antibody among those studied in clinical trials. In addition, treatment with C04 caused relatively less body weight reduction than CP-870,893 treatment, suggesting that C04 had a favorable toxicity profile (Fig. 6D).

## DISCUSSION

Here we described a droplet-based microfluidics platform for functional screening of millions of antibodies. The platform shared some key features with the most efficient selection methods such as phage display *(8–13)*. First, the genotype and phenotype linkage was maintained through the whole process. And second, the product from one round can be directly amplified and used as the input of the next round of selection. Thus multiple rounds of iteration allowed enrichment of rare hits. Comparing to the conventional method to individually express and assay thousands of antibodies, the throughput of this platform increased to 10 millions. It is especially useful in the development of next generation cancer immunotherapies, such as agonist antibodies or bispecific antibodies when simple binding assay maybe inadequate.

In order to demonstrate the usefulness of the platform, we first applied the platform to discover bispecific antibody and agonist antibody, whose development was limited by low diversity and/or low throughput and potentially biased screening.

Bispecific T or NK cell engager (BiTE or BiTE) also hold great promise for cancer treatment and a growing number of BiTE and BiTE are making their way through various stages of development *(6)*. To obtain the optimal BiTE or BiTE, a bispecific antibody library is constructed to cover the complexity of the array of tumor antigen targeting antibodies. The difficulties arise, however, with the large number of bispecific antibodies in the library exceeding the throughput of the existing methods notwithstanding the approach described herein provides significant opportunities to screen large bispecific antibody library.

The areas of costimulatory receptor agonist has been reignited over the last decade thanks to the substantial advances in the field of immunoncology. The costimulatory receptors are expressed on a number of immune cell types, including T cells, B cells and natural killer (NK) cells as well as APCs, and engagement of these receptors can promote immune cell function, proliferation and survival. Nevertheless, there are no general rules to guide the screening of agonist antibody. For example, a panel of antibodies bind to the same or similar epitopes of Fas receptor but result in different biological effects, with some acting as agonists and others as antagonists *(4, 26)*. The intrinsic complexity of agonist antibody required screening as many antibodies as possible. When we used the unique platform to screen CD40 agonist antibody, we found a few potent CD40 agonist antibodies, most of which were too rare (<0.02% frequency) to be discovered by using the conventional method.

The method can also be applied to high throughput analysis of cell-cell interactions. For example, we can envision a scenario where DC cells infected with lentivirus library encoding neoantigens are coencapsulated with tumor infiltrating T cells to map the pairs of cognate antigens and T cell receptors (TCR) *(27)*. The method can also be adapted to screen different types of molecules such as cytokines *(28–30)*. The innovative applications of this activity based selection method have been limited only by the imagination of the users. Altogether, we developed a high-throughput general platform for function-based screening up to millions of antibodies. With the throughput in the millions of antibody producing cells and without any pre-assumptions other than the key function used for screening, this may revolutionize the next generation cancer immunotherapy drug development, as well as basic researches involving cell-to-cell interactions.

## MATERIAL AND METHODS

### Cell culture

HEK293FT cells (Thermo Fisher Scientific, R70007) were cultured in DMEM medium (Thermo Fisher Scientific). Jurkat cells (ATCC, TIB-152) were cultured in RPMI 1640 medium (Thermo Fisher Scientific). SKBR-3 cells (ATCC, HTB-30) were cultured in McCoy’s 5A (Modified) Medium (Biological Industries). All the culture medium was supplemented with 10% fetal bovine serum (Biological Industries), 1× non-essential amino acids, 100 U/ml penicillin, 100 μg/ml streptomycin and 12.5mM HEPES (Thermo Fisher Scientific). HEK293F cells (Thermo Fisher Scientific, R79007) were suspension cultured in FreeStyle™ 293 Expression Medium (Thermo Fisher Scientific). All the cells were maintained in CO2 incubator at 37°C.

### IL2 reporter cell line

Jurkat/pIL2-eGFP was developed to monitor activation of T cells. The Jurkat cell line was transfected with vector carrying eGFP reporter gene under control of full-length IL-2 promoter (the region from −648 to −1 upstream of translation initiation codon of IL2). After stimulated by anti-CD3 and anti-CD28 antibodies, cells expressing high level of GFP were sorted into wells of 96 well plate using FACS and individual clones of Jurkat/pIL2-eGFP cell line were characterized.

### CD40 reporter cell line

Jurkat/NF-κB-GFP-hCD40 reporter cell was developed to monitor activation of CD40. The Jurkat cell was transfected with vector carrying eGFP reporter gene under control of NF-κB response element. After stimulated by 10 ng/mL TNFα (Sino Biological), cells expressing high level of eGFP were sorted by FACS and the Jurkat/NF-κB-eGFP cell line were characterized. The resulting Jurkat/NF-κB-eGFP cell line was infected with lentivirus expressing full-length human CD40. After stimulated by 100 nM hexameric CD40L Fc fusion protein, individual cells with high level of GFP signal were sorted into wells of 96 well plate by Fluorescence Activated Cell Sorter (FACS) and individual clones of Jurkat/NF-κB-GFP-hCD40 cell were characterized.

### Microfluidic chip fabrication

All microfluidic chips were fabricated in polydimethylsiloxane (PDMS) polymer (Sylgard 184 elastomer kit; Dow Corning Corp) using the standard soft lithography as described *(31)*. Masters were made using one layer of SU-8 photoresist (MicroChem). The depth of the two devices is 40+/-1μm to allow the droplet generating or flowing in a monolayer format. For device ii, the PDMS is bonded to a piezoelectric substrate (Y128-cut Lithium niobate wafer) where a golden interdigital electrode is patterned with standard lift-off technology and aligned with the fluidic channel above. Microfluidics devices were treated before use with 1% v/v 1H,1H,2H,2H-perfluorodecyltrichlorosilane (Alfa Aesar) in Novec HFE7500 fluorinated oil (3M) to prevent droplets wetting the channel walls.

### Phage display

A human naïve scFv library was constructed from PBMC of 30 healthy donors with standard protocols. The phage library was incubated with biotinylated CD40-Fc fusion protein or Her2 recombinant protein (Acro biosystems) for 2 hours at RT and the phage-antigen complex was captured by Dynabeads M280 (Life technologies). The bound phages were eluted by Glycin-HCl (pH 2.2) for 10 min at RT and neutralized with Tris-HCl (pH 8.0) to adjust pH to 7.5. The phagemid DNA was isolated using plasmid miniprep kit (Qiagen).

### Lentivirus library construction

Both phagemids and lentiviral vector pLV-ef1α-ScFv-Fc were digested with enzyme SfiI. The lentiviral vector and scFv genes were isolated after electrophoresis. ScFv genes were ligated into lentiviral vector. The product of ligation reaction were transformed into XL1-Blue competent cells by electroporation transformation technique and most of the transformed bacteria were plated on LB Agar plate. The remaining bacteria were serial diluted and plated to estimate the size of library. The lentiviral plasmid was prepared using plasmid midiprep kit (Qiagen) for lentivirus preparation.

### Lentivirus preparation

When cell confluency reached 80%, HEK293FT cells were transfected with lentiviral backbone plasmid and packaging plasmids using PEI transfection reagent. The medium was changed to fresh complete culture medium 6 h after transfection. Supernatant containing lentivirus were harvested after 48 h, centrifuged at 300g for 5 min at 4°C and filtered by 0.45μm filter to remove cell debris. The titer of virus was measured using P24 ELISA kit (Clontech).The virus was aliquoted and stored at −80°C.

### Function based screening of anti-Her2/anti-CD3 bispecific antibody using microfluidics

Aqueous phase I: Jurkat/NF-κB-eGFP reporter cells were washed with PBS and stained with 1μM CellTrace Yellow dye for 10 min at 37°C. The stained Jurkat/NF-κB-eGFP reporter cells were washed 2 times with RPMI 1640 and resuspended in cell culture medium (RPMI 1640, 5%FBS, 25mM HEPES and 0.1%Pluronic F-68) containing 1 μg/mL anti-CD28 antibody (Invitrogen).

Aqueous phase II: The stable Her2 expressing K562 cells were infected with lentiviral antibody library. The resulting antibody secreting K562 were washed with PBS and stained with 1μM CellTrace Violet dye for 10 min at 37 °C. The stained cells were washed 2 times with RPMI 1640 and resuspended with cell culture medium containing 200nM DY647. K562 cells infected with positive control lentivirus were resuspended with cell culture medium containing 1500nM DY647.

The Aqueous phase I and II were injected into the droplet generation chip from different inlets and used as the disperse phase. Novec HFE7500 fluorinated oil (3M) containing 2% w/w fluoro-surfactant (RAN Biotechnologies) was used as continue phase to produce droplets with average size of 100pL. The flow rates of aqueous phase I, aqueous phase II and oil phase were adjusted so on average 1 reporter cell and 0.5 antibody secreting cell were coencapsulated per droplet. During droplet production, the cell suspension was cooled using ice-water to inhibit antibody secretion. The droplets were collected and incubated at 37°C for 16 h.

The droplets were first gated to eliminate coalesced droplets and retain only droplets of desired size. Positive control and the screening population droplets were distinguished based on the different intensity of fluorescence of DY647. For the screening population, the droplets were selected for the presence of Jurkat reporter cells based on CellTrace Yellow signal and K562 cells based on CellTrace Violet signal. The FADS was performed to sort the droplets containing Jurkat emitting GFP fluorescence. Finally we gated and sorted the droplets with GFP signal colocalized with Jurkat staining signal, but not with K562 signal.

The cells were recovered from the sorted droplets by adding 200μL RPMI 1640 medium supplemented with 10% Fetal Bovine Serum (FBS) and 24% Nycodenz, followed adding 50μL 1H,1H,2H,2H-Perfluoro-1-octanol (Sigma). After mixing thoroughly and centrifuged at 300g for 5 min at 4°C, the aqueous layer was completely separated and washed with RPMI 1640 medium. The recovered cells were lysed and antibody genes were amplified from the cells.

### Anti-Her2/anti-CD3 bispecific antibody in vitro experiment

#### Jurkar/IL-2-GFP reporter cell assay

For detecting the activity of anti-Her2/anti-CD3 antibody candidates, Jurkat/IL-2-GFP reporter cells were stimulated with different concentrations of anti-Her2/anti-CD3 antibodies and 1μg/ml anti-CD28 antibody in presence of K562 cells or K562-Her2 cells. After 16h incubation, GFP expression of reporter cell was detected by flow cytometry.

#### Anti-Her2/anti-CD3 bispecific antibody ex vivo assay

Primary T cells were isolated from PBMCs using CD3 MicroBeads (Miltenyi, 130-050-101). 1×10^5^ T cells were cocultured with target cells (SKBR3 cells, MDA-MB-231 cells or HEK293 cells) at a 1:1 ratio. Different concentrations of anti-Her2/anti-CD3 antibody or control antibody together with 1μg/ml anti-CD28 antibody were added. After 48 h incubation, cells were collected and stained with anti-CD3-FITC (BioLengend, 300406) and anti-CD69-APC (BioLegend, 310910) for 30 min at 4 °C. T cell activation was determined by flow cytometry. Flow cytometry results were analyzed with software Flowjo X.

Cell supernatant was collected to quantify the cytokine release and cytotoxicity. IL-2 and INF-γ were measured with ELISA kit according to the manufacturer’s instructions. Cytotoxicity was analyzed by measuring levels of released lactate dehydrogenase (LDH) using the CytoTox 96 non-radioactive cytotoxicity assay protocol (Promega).

#### Function based screening of CD40 agonist antibody using microfluidics

Aqueous phase I: Jurkat-CD40-NFκB reporter cells were washed with PBS and stained with 1μM CellTrace Yellow dye for 10 min at 37°C. The stained Jurkat-CD40-NFκB reporter cells were washed 2 times with DMEM and resuspended at 20 million cells/mL with cell culture medium (DMEM, 5%FBS, 25mM HEPES and 0.1%Pluronic F-68) containing 16.67nM Dylight647 conjugated goat anti-human Fc IgG and 24% Nycodenz.

Aqueous phase II: For the screening population, the HEK293FT cells were infected with lentiviral antibody library and resuspended with cell culture medium containing 500nM DY405. For positive control droplets, HEK293FT cells were resuspended with cell culture medium containing soluble hexameric CD40L protein and 1500nM DY405. For negative control droplets, HEK293FT cells were resuspended with cell culture medium containing ani-HEL antibody and 2500nM DY405.

The aqueous phase I and II were injected into the droplet generation chip from different inlets and used as the disperse phase. Novec HFE7500 fluorinated oil (3M) containing 2% w/w fluoro-surfactant (RAN Biotechnologies) was used as continue phase to produce droplets with average size of 100pL. The flow rates of aqueous phase I, aqueous phase II and oil phase were adjusted so on average 1 reporter cell and 0.5 antibody secreting cell were coencapsulated per droplet. During droplet production, the cell suspension was cooled using ice-water to inhibit antibody secretion. The droplets were collected and incubated at 37°C for 16 h.

The droplets were first gated to eliminate coalesced droplets and retain only droplets of desired size. Negative control droplets, positive control droplets and screening droplets were distinguished based on the different intensity of blue fluorescence of DY405. For screening population, the droplets were selected for the presence of Jurkat reporter cells in the droplet based on the yellow fluorescence of CellTrace Yellow dye. Finally the FADS was performed to sort the droplets containing Jurkat emitting fluorescence of Dylight647 and GFP. The cells were recovered from the sorted droplets by adding 200μL DMEM medium supplemented with 10%FBS and 24% Nycodenz, followed adding 50μL 1H,1H,2H,2H-Perfluoro-1-octanol (370533, Sigma). After mixing thoroughly and centrifuged at 300g for 5min at 4°C, the aqueous layer was completely separated and washed with DMEM medium. The cells were resuspended with DMEM medium containing 10% FBS and 1% PS (Penicillin Streptomycin) and then cultured for 1-2 weeks.

#### Bioinformatic analysis of the PacBio sequencing results

Single molecule real-time (SMRT) sequencing platform (Pacific Biosciences) generated long sequencing read with an average read length of ∼20 kb, which could sequence a scFv (subreads) more than 10 times. Circular consensus sequencing program from PacBio SMRTportal software (version 4.1.0) can take multiple subreads of the same SMRTbell sequence and combines them, employing a statistical model, to produce one high quality circular consensus sequence (CCS). After CCS from long sequencing reads were generated, scFv flanking sequences were trimmed, scFv DNA sequences were translated into protein using CLC genomics workbench (version 11.0.1) and CDR1-3 regions for heavy and light chains were identified by IgBLAST (version 1.15.0). CD-hit (version 4.8.1) was used to group scFvs at protein similarity 95%. Frequency of scFvs for each round were calculated in java program, respectively. ScFvs that appeared in only one sample were removed because those are likely PCR artifact product. The barplot were draw by R packages ggplot2 (version 3.2.1).

#### Protein expression and purification of fill length IgG

Equal amounts of heavy chain and light chain expression plasmids were co-transfected into 293F cells. Five days after transfection, the cells were centrifuged at 3000rpm for 10min at 4°C and the supernatants were harvested and passed through a 0.45μm filter. Antibodies were purified with HiTrap Protein A column (GE) using ÄKTA purifier chromatography system.

### CD40 in vitro experiment

#### Species Cross Reactivity of antibody

Cross-reactivity of C03 was assessed by flow cytometry analysis. Briefly, HEK293FT cells were transiently transfected with human or rhesus macaque CD40 expressing vector using PEI (polyscience). After 48 hours HEK293FT-hCD40 or HEK293FT-rCD40 cells were incubated with different concentrations of antibody at RT for 30 min. Then the cells were stained with AlexFuor488-conjugated goat anti human Fc (Life technologies) at RT for 30 min and analyzed by flow cytometry. Fluorescence intensity is equal to the percentage of GFP positive cells multiplied by Median Fluorescence Intensity (MFI). The fluorescence intensity was plotted against the antibody concentrations using software GraphPad Prism.

#### Surface plasmon resonance (SPR) analysis

SPR experiments were performed with a Biacore T200 SPR system (GE Healthcare). In brief, experiments were performed at 20°C in HBS-P+ buffer (0.01 M HEPES, 0.15 M NaCl and 0.05% v/v Surfactant P20). Anti-his antibody was immobilized on Series S CM5 chip by amine coupling, his-tagged cynomolgus monkey CD40 were captured by the immobilized anti-his antibody with a flow rate of 10μL min−1 for 60s. Twofold serially diluted CD40 antibodies were injected through flow cells for 120 s followed by a 130 s dissociation phase at a flow rate of 30 μL min−1. Prior to next assay cycle, the sensor surface was regenerated with Glycine-HCl (pH 1.5) for 30 s at a flow rate of 30 μL min−1. Background binding to blank immobilized flow cells was subtracted, and KD values were calculated using the 1:1 binding kinetics model built in the BIAcore T200 Evaluation Software (version 3.2).

#### CD40 antibody selectivity

Human CD40 (Acrobiosystems), GITR (Acrobiosystems), OX40 (Acrobiosystems), 4-1BB (Acrobiosystems) or BSA (Solarbio) were plated onto a microtiter plate at 4°C overnight. The coated wells were blocked by 0.5% BSA in PBS at 37°C for 1 h. Serial diluted antibodies were added and incubated at 37°C for 1h, washed 8 times, goat anti-human IgG-HRP (SouthernBiotech) was added and incubated at 37°C for 30 min. After 8 times washes, ABTS substrate solution (Thermofisher) was added and the OD at 405 nm were measured with a plate reader.

#### Jurkat/NF-κB-GFP-hCD40 reporter cell assays

For CD40 agonists activity detection, Jurkat/NF-κB-GFP-hCD40 reporter cells were incubated with different concentrations of CD40 agonist antibodies with or without goat anti-human Fc antibody (SouthernBiotech) for 24 hours. GFP expression was detected by flow cytometry.

For FcγRIIB dependency experiment, HEK293FT cells were transiently transfected with FcγRIIB expressing vector using PEI. After 36 hours HEK293FT-FcγRIIB cells were plated in 48-well plate and cultured overnight at 37°C.Then Jurkat/NF-κB-GFP-hCD40 reporter cells and different concentrations of C03 or HEL were cocultured with HEK293FT-FcγRIIB cells for 24 hours. GFP expression was detected by flow cytometry.

For the data analysis of Jurkat/NF-κB-GFP-hCD40 reporter cell assays, flow cytometry results were analyzed with software Flowjo X, fluorescence intensity is equal to the percentage of GFP positive cells mutiplied by MFI. The fluorescence intensity of cells was plotted against the antibody concentrations was calculated using software GraphPad Prism.

#### CD40 ex vivo experiment

Following thawing and recovery of human PBMCs, monocytes were selected by adhering to plastic and then cultured for 8 days in RPMI containing 10% FBS (Gibico), 100 ng/mL GM-CSF (R&D Systems) and 10 ng/mL IL-4 (R&D Systems). Suspended cells were harvested and confirmed to be dendritic cells by CD11c expression. B cells were isolated from PBMCs by magnetic selection using CD19 beads (Miltenyi). 1 × 10^5^ dendritic cells or B cells were incubated with different concentrations of C03 with or without goat anti-human Fc antibody (SouthernBiotech) for 48 hours. Upregulation of the activation markers CD86 was analyzed by flow cytometry (Biolegend). Flow cytometry results were analyzed with software Flowjo X. The MFI of cells was plotted against the antibody concentrations using software GraphPad Prism.

#### CD40 in vivo experiment

FcγR/CD40-humanized mice have been described previously and were kindly provided by Dr Jeffrey Ravetch (The Rockefeller University). Mice were bred and maintained in the specific-pathogen-free environment at the Department of Laboratory of Animal Science, Shanghai Jiao Tong University School of Medicine. All animal care and study were performed in compliance with institutional and NIH guidelines and had been approved by SJTUSM Institutional Animal Care and Use Committee (Protocol Registry Number: A-2015-014).

#### OVA-specific CD8+ T-cell response model

CD40/FcγR humanized mice were adoptively transferred with CD45.1+ splenic OT-I cells (2 × 10^6^ cells in 200 μl PBS per mouse) via tail vein injection one day before immunized with 2 μg of DEC-OVA, in the presence of the CD40 agonist antibody or the isotype control by intraperitoneal injection. On day 6, spleen cells were harvested. After red blood cells lysis, the single-cell suspension was stained with anti-CD4 (clone RM4-5), anti-CD8 (clone 53-6.7), anti-CD45.1 (A20), anti-TCR-Vα2 (B20.1) to quantify OVA-specific OT-I CD8+ T cells. OT-I CD8+ T cell is defined as CD45.1+CD8+TCR-Vα2+ cells.

#### Syngeneic mouse model

CD40/FcγR humanized mice were inoculated subcutaneously with 2×10^6^ MC38 cells. When tumor volumes reached 50 to 100 mm^3^, mice were randomly assigned to different groups (n=5). MC38-bearing CD40/FcγR humanized mice were treated intraperitoneally with C03, CP870893 or HEL (3 mg/kg, q3d×2). Tumor growth was monitored every 3 days by measuring the length (L) and width (W) with calipers and tumor volume was calculated with the formula, (L×W^2^)/2.

## Acknowledgments

This work was supported by the National Natural Science Foundation of China (grant numbers 81872787), China Postdoctoral Science Foundation (grant number 2020M670625), the Fundamental Research Funds for the Central Universities, Nankai University(grant number ZB19100123 and 63191212), Natural Science Foundation of Tianjin (grant number 19JCZDJC32900), Key Laboratory of Immune Microenvironment and Disease open funding (grant number 20180102), Shanghai Municipal Science and Technology Commission (project No. 19431902900), National key Research and Development plan of China (grant number 2018YFE0200401 and 2017YFA0504801).F.L and Y.Z are also supported by the innovative research team of high-level local universities in Shanghai (SSMU-2DCX20180100).The work was also sponsored by HiFiBio Therapeutic funding.

## Author Contributions

H.Z. designed all the experiments. Y.W., R.J., B.S., W.W., Y.Z., M.H., P.F., S.W., P.M., N.L., R.W., P.M. and R.L. performed the experiments. H.Z., Y.W., B.S., H.Z., L.S., F.L. and Y.C analyzed all the data and wrote the manuscript with the input of all authors.

## Competing interests

The authors have declared that no competing interest exists.

## Notes

### Competing Interest Statement

The authors have declared no competing interest.

## REFERENCES

1. L. P. Andrews, H. Yano, D. A. A. Vignali, Inhibitory receptors and ligands beyond PD-1, PD-L1 and CTLA-4: breakthroughs or backups. Nat. Immunol. 20, 1425–1434 (2019).

2. J. Tang, J. X. Yu, V. M. Hubbard-Lucey, S. T. Neftelinov, J. P. Hodge, Y. Lin, Trial watch: The clinical trial landscape for PD1/PDL1 immune checkpoint inhibitors. Nat. Rev. Drug Discov. 17, 854–855 (2018).

3. A. Hoos, Development of immuno-oncology drugs - from CTLA4 to PD1 to the next generations. Nat. Rev. Drug Discov. 15, 235–247 (2016).

4. P. A. Mayes, K. W. Hance, A. Hoos, The promise and challenges of immune agonist antibody development in cancer. Nat. Rev. Drug Discov. 17, 509–527 (2018).

5. R. H. Vonderheide, CD40 Agonist Antibodies in Cancer Immunotherapy. Annu. Rev. Med. 71, 47–58 (2020).

6. A. F. Labrijn, M. L. Janmaat, J. M. Reichert, P. Parren, Bispecific antibodies: a mechanistic review of the pipeline. Nat. Rev. Drug Discov. 18, 585–608 (2019).

7. H. Li, P. Er Saw, E. Song, Challenges and strategies for next-generation bispecific antibody-based antitumor therapeutics. Cell. Mol. Immunol. 17, 451–461 (2020).

8. G. P. Smith, Phage Display: Simple Evolution in a Petri Dish (Nobel Lecture). Angew. Chem. Int. Ed. Engl. 58, 14428–14437 (2019).

9. R. A. Lerner, Manufacturing immunity to disease in a test tube: the magic bullet realized. Angew. Chem. Int. Ed. Engl. 45, 8106–8125 (2006).

10. A. R. Bradbury, S. Sidhu, S. Dubel, J. McCafferty, Beyond natural antibodies: the power of in vitro display technologies. Nat. Biotechnol. 29, 245–254 (2011).

11. P. Amstutz, P. Forrer, C. Zahnd, A. Pluckthun, In vitro display technologies: novel developments and applications. Curr. Opin. Biotechnol. 12, 400–405 (2001).

12. E. T. Boder, K. D. Wittrup, Yeast surface display for screening combinatorial polypeptide libraries. Nat. Biotechnol. 15, 553–557 (1997).

13. G. Georgiou, C. Stathopoulos, P. S. Daugherty, A. R. Nayak, B. L. Iverson, R. C. Iii, Display of heterologous proteins on the surface of microorganisms: From the screening of combinatorial libraries to live recombinant vaccines. Nat. Biotechnol. 15, 29–34 (1997).

14. S.L. Anna., N. Bontoux., H. A. Stone., Formation of dispersions using “flow focusing” in microchannels. Applied Physics Letters 82, 364–366 (2003).

15. L. Mazutis, J. Gilbert, W. L. Ung, D. A. Weitz, A. D. Griffiths, J. A. Heyman, Single-cell analysis and sorting using droplet-based microfluidics. Nat. Protoc. 8, 870–891 (2013).

16. N. Shembekar, C. Chaipan, R. Utharala, C. A. Merten, Droplet-based microfluidics in drug discovery, transcriptomics and high-throughput molecular genetics. Lab Chip 16, 1314–1331 (2016).

17. G. Georgiou, G. C. Ippolito, J. Beausang, C. E. Busse, H. Wardemann, S. R. Quake, The promise and challenge of high-throughput sequencing of the antibody repertoire. Nat. Biotechnol. 32, 158–168 (2014).

18. B. El Debs, R. Utharala, I. V. Balyasnikova, A. D. Griffiths, C. A. Merten, Functional single-cell hybridoma screening using droplet-based microfluidics. Proc. Natl. Acad. Sci. U. S. A. 109, 11570–11575 (2012).

19. N. Shembekar, H. Hu, D. Eustace, C. A. Merten, Single-Cell Droplet Microfluidic Screening for Antibodies Specifically Binding to Target Cells. Cell Rep. 22, 2206–2215 (2018).

20. K. Eyer, R. C. L. Doineau, C. E. Castrillon, L. Briseño-Roa, Single-cell deep phenotyping of IgG-secreting cells for high-resolution immune monitoring. Nat. Biotechnol. 35, 977–982 (2017).

21. A. Gérard, A. Woolfe, G. Mottet, M. Reichen, C. Castrillon, High-throughput single-cell activity-based screening and sequencing of antibodies using droplet microfluidics. Nat. Biotechnol. 38, 715–721 (2020).

22. H. Zhang, E. Sturchler, J. Zhu, A. Nieto, P. A. Cistrone, J. Xie, L. He, K. Yea, T. Jones, R. Turn, P. S. Di Stefano, P. R. Griffin, P. E. Dawson, P. H. McDonald, R. A. Lerner, Autocrine selection of a GLP-1R G-protein biased agonist with potent antidiabetic effects. Nat Commun 6, 8918 (2015).

23. H. Zhang, I. A. Wilson, R. A. Lerner, Selection of antibodies that regulate phenotype from intracellular combinatorial antibody libraries. Proc. Natl. Acad. Sci. U. S. A. 109, 15728–15733 (2012).

24. L. Schmid, D. A. Weitz, T. Franke, Sorting drops and cells with acoustics: acoustic microfluidic fluorescence-activated cell sorter. Lab Chip 14, 3710–3718 (2014).

25. R. Lutterbuese, T. Raum, R. Kischel, P. Hoffmann, S. Mangold, B. Rattel, M. Friedrich, O. Thomas, G. Lorenczewski, D. Rau, E. Schaller, I. Herrmann, A. Wolf, T. Urbig, P. A. Baeuerle, P. Kufer, T cell-engaging BiTE antibodies specific for EGFR potently eliminate KRAS- and BRAF-mutated colorectal cancer cells. Proc. Natl. Acad. Sci. U. S. A. 107, 12605–12610 (2010).

26. M. Chodorge, S. Zuger, C. Stirnimann, C. Briand, L. Jermutus, M. G. Grutter, R. R. Minter, A series of Fas receptor agonist antibodies that demonstrate an inverse correlation between affinity and potency. Cell Death Differ 19, 1187–1195 (2012).

27. T. Kula, M. H. Dezfulian, C. I. Wang, N. S. Abdelfattah, Z. C. Hartman, K. W. Wucherpfennig, H. K. Lyerly, S. J. Elledge, T-Scan: A Genome-wide Method for the Systematic Discovery of T Cell Epitopes. Cell 178, 1016–1028 e1013 (2019).

28. J. B. Spangler, I. Moraga, J. L. Mendoza, K. C. Garcia, Insights into cytokine-receptor interactions from cytokine engineering. Annu. Rev. Immunol. 33, 139–167 (2015).

29. J. T. Sockolosky, E. Trotta, G. Parisi, L. Picton, L. L. Su, A. C. Le, A. Chhabra, S. L. Silveria, B. M. George, I. C. King, M. R. Tiffany, K. Jude, L. V. Sibener, D. Baker, J. A. Shizuru, A. Ribas, J. A. Bluestone, K. C. Garcia, Selective targeting of engineered T cells using orthogonal IL-2 cytokine-receptor complexes. Science 359, 1037–1042 (2018).

30. P. N. Kelly, Engineering cytokine-receptor pairs. Science 359, 1004 (2018).

31. J. C. McDonald, G. M. Whitesides, Poly(dimethylsiloxane) as a material for fabricating microfluidic devices. Acc. Chem. Res. 35, 491–499 (2002).

