## Supplementary Information for "High throughput functional screening for next generation cancer immunotherapy using droplet-based microfluidics"

**Supplementary Materials**

Fig.S1

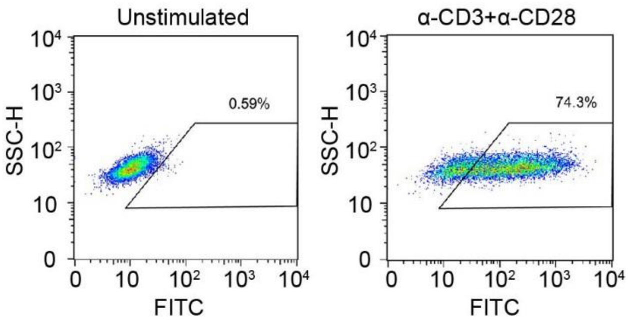


**Fig. S1. Development and validation of Jurkat/pIL2-eGFP reporter cells.** The Jurkat/pIL2-eGFP reporter cells were stimulated with anti-CD3 and anti-CD28 antibody overnight. GFP fluorescence was analyzed by flow cytometry.

Fig.S2

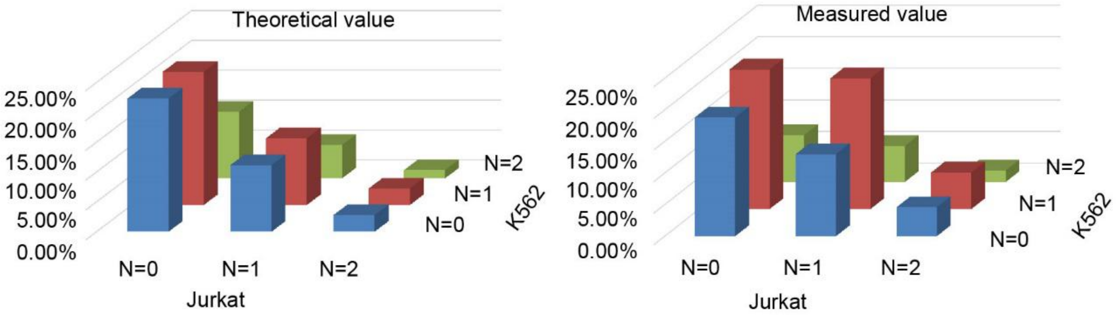


**Fig. S2. The double Poisson distribution of cell number in droplets.** K562-Her2 and Jurkat reporter cells were stained with CellTrace Violet and CellTrace Yellow,respectively. Cell loading was evaluated by counting the cell labeling signals.

Fig.S3

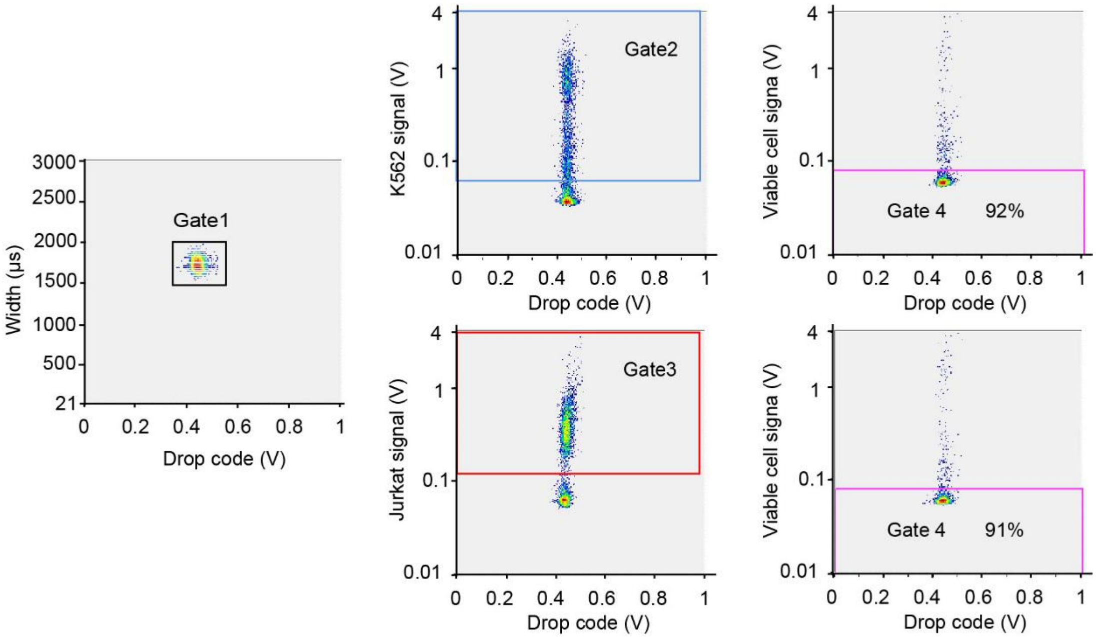


**Fig. S3. The gating strategy for the analysis of cell viability.** The droplets were first gated (gate 1) to eliminate coalesced droplets and retain only droplets of the desired size. The droplet in gate 2 defines droplet containing K562-Her2 cells; gate 3 defines the droplet containing Jurkat cells. Gate 4 defines the viable cells with low fluorescent nuclear staining, indicating the live cells after 16 h incubation.

Fig.S4

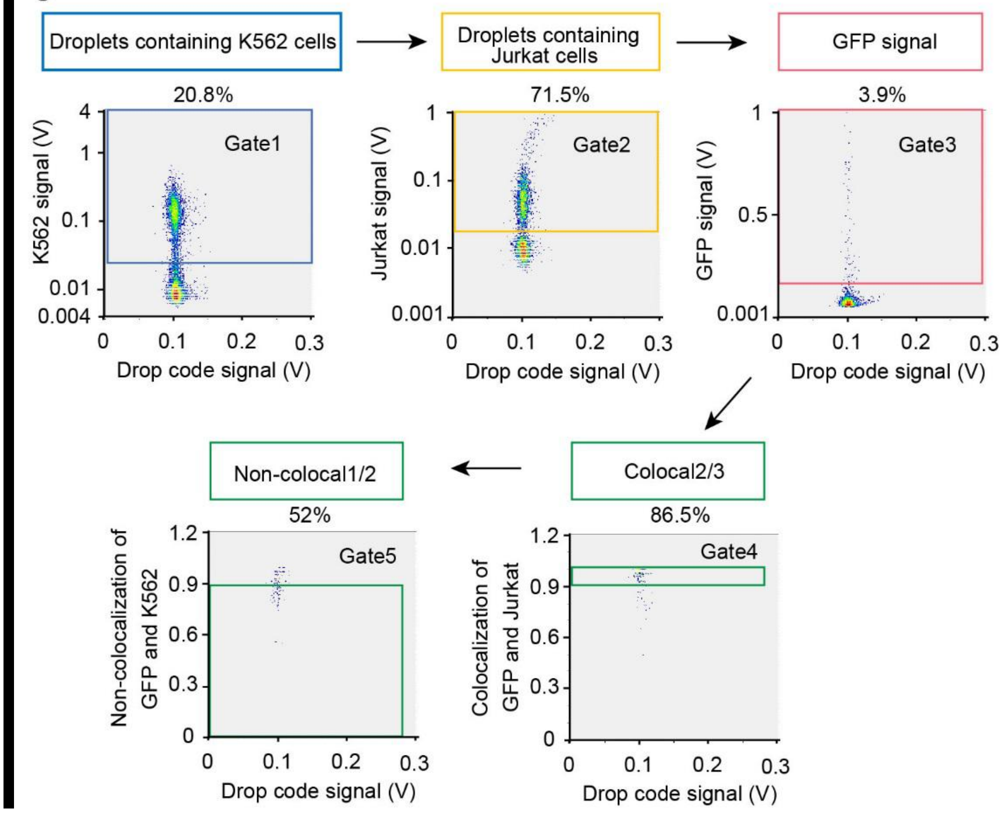


**Fig. S4. The gating strategies for screening anti-Her2/anti-CD3 bispecific antibody.** The droplets were first gated to eliminate coalesced droplets and retain only droplets of the desired size. The droplets containing K562 cell were gated based on CellTrace Violet fluorescence signal (gate 1). CellTrace Yellow fluorescence signal peak showed the presence of reporter cell in the droplet (gate 2). GFP fluorescence signal peak indicated activation of the reporter cell (gate 3). Lastly colocalization2/3 and non-colocalization1/2 were used to sort droplets where GFP was from Jurkat rather than K562 (gate 4 and gate 5).

Fig.S5

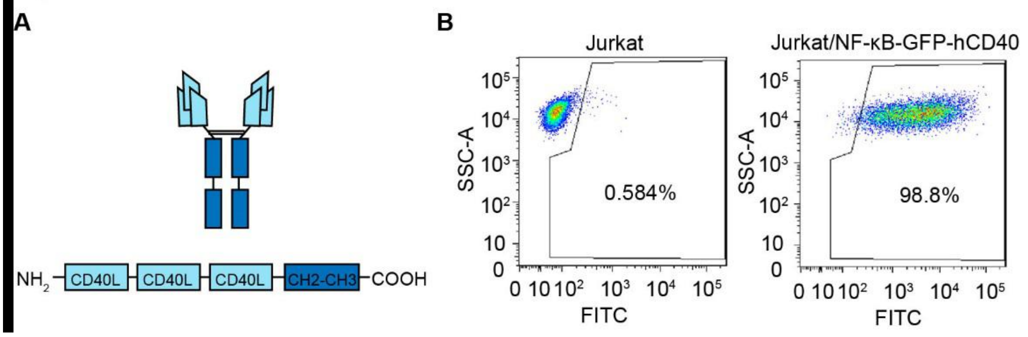


**Fig. S5. Development and validation of Jurkat/NF-κB-GFP-hCD40 reporter cells.** (A) Diagram of the hexameric CD40L. Three receptor binding domains of CD40 ligand were tandemly linked to form a trivalent protein. IgG1-Fc is then used to link two of the trivalent proteins together, creating six receptor binding domains in a single agonist. (B) The Jurkat/NF-κB-GFP-hCD40 reporter cells were stimulated with hexameric form of CD40L overnight. GFP fluorescence was analyzed using flow cytometry.

Fig.S6

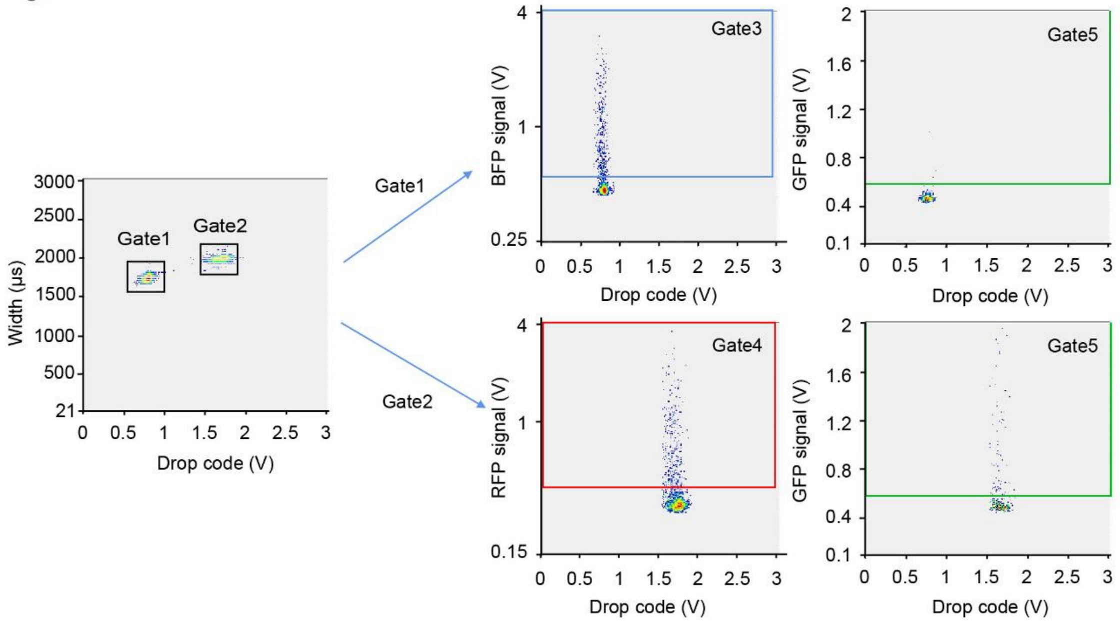


**Fig. S6. The gating strategy for the analysis of Jurkat/NF-κB-GFP-hCD40 activation in droplet.** The droplets were first gated to eliminate coalesced droplets and retain only droplets of the desired size. Two gates were assigned using drop code DY638. Gate 1 defines droplets from negative emulsion where HEL cells were encapsulated, gate 2 defines droplets from positive emulsion where CD40L cells were encapsulated. Gate 3 defines the droplet containing HEL cells according to BFP signal and gate 4 defines the CD40L cells according to RFP signal. Gate 5 defines droplets where reporter cells were activated and emitted GFP fluorescence. After 16 h incubation, 24% of the CD40L cell containing droplets exhibited GFP fluorescence signal while the HEL cell and reporter cell coencapsulating droplets showed clean background of the activation of reporter cell (0.5%).

**Fig.S7**

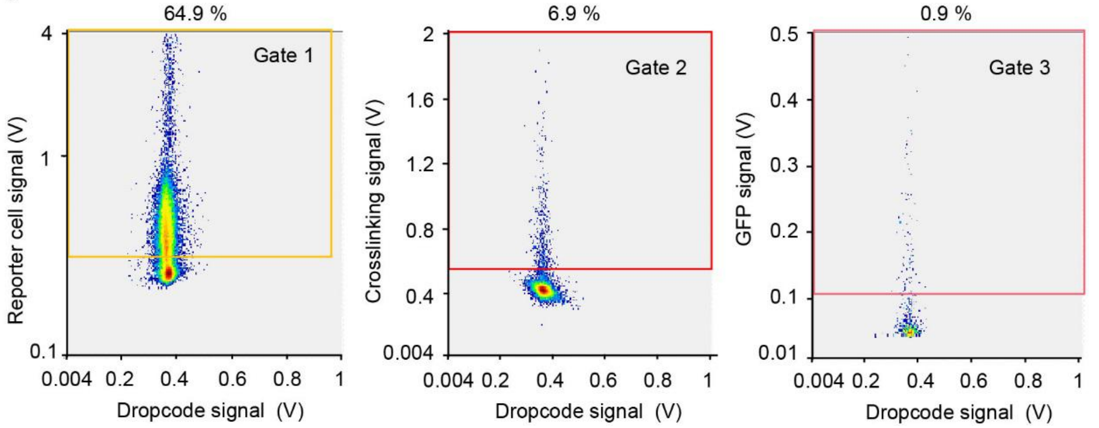


**Fig. S7**. **The gating strategies for screening CD40 agonist antibody.** The droplets were first gated to eliminate coalesced droplets and retain only droplets of the desired size. The droplets of the screening population were first gated based on the intensity of DY405. CellTrace Yellow fluorescence signal peak showed the presence of reporter cell in the droplet (gate 2). The Dylight647 fluorescence peak signal indicated binding of the secreting antibodies to CD40 on the reporter cell (gate 3) and GFP fluorescence signal peak indicated activation of the reporter cell (gate 4).

**Fig.S8**

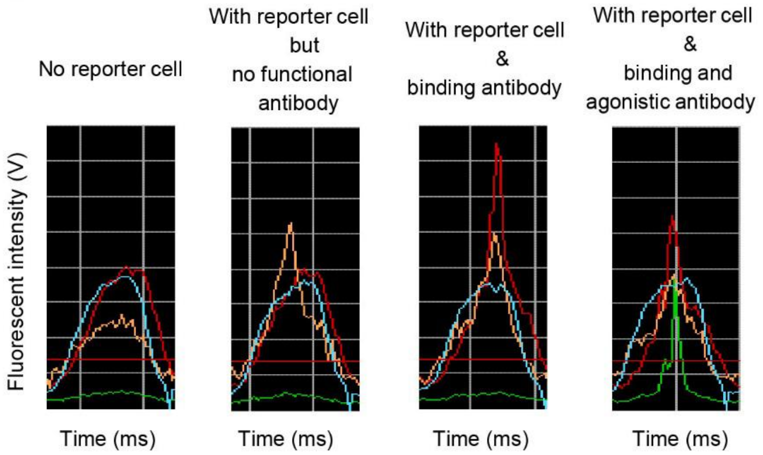


**Fig. S8**. **Experimental time traces corresponding to the examples in Fig. 6A.** Droplets are sorted if they display a green peak (green line, GFP signal), a red peak (red line, rabbit anti-human IgG Fc DL650 gathering around reporter cells) and a yellow peak (yellow line, reporter cells) that co-localize. Blue fluorescence (blue line) signal is used to identify droplets population.

Fig.S9

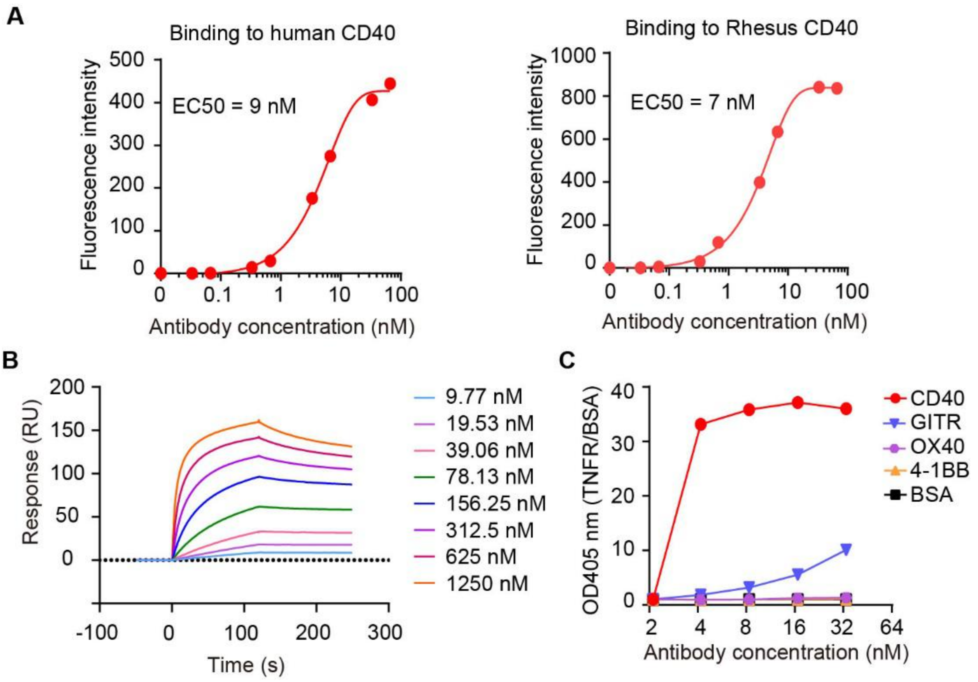


**Fig. S9. Binding of C04 to human,rhesus and cynomolgus monkey CD40.** (A) Binding of C04 to human and rhesus CD40 was determined by flow cytometry. 293T cells were transient transfected with human or rhesus CD40 and incubated with different concentration of antibody C04 and goat anti-human IgG Fc Alexa Fluor 488. The cells were analyzed by flow cytometry. (B) Binding of C04 to cynomolgus monkey CD40 was determined by SPR analysis. Anti-his antibody was immobilized on Series S CM5 chip, his-tagged cynomolgus monkey CD40 were captured by the immobilized anti-his antibody. Different concentrations of CD40 antibodies were injected through flow cells and Kd values were calculated using the 1:1 binding kinetics model. (C) Binding of C04 to different tumor necrosis factor superfamily (TNFSF) receptors was determined by ELISA.
